# Riemannian geometry and statistical modeling correct for batch effects and control false discoveries in single-cell surface protein count data from CITE-seq

**DOI:** 10.1101/2020.04.28.067306

**Authors:** Shuyi Zhang, Jacob R. Leistico, Christopher Cook, Yale Liu, Raymond J. Cho, Jeffrey B. Cheng, Jun S. Song

## Abstract

Recent advances in next generation sequencing-based single-cell technologies have allowed high-throughput quantitative detection of cell-surface proteins along with the transcriptome in individual cells, extending our understanding of the heterogeneity of cell populations in diverse tissues that are in different diseased states or under different experimental conditions. Count data of surface proteins from the cellular indexing of transcriptomes and epitopes by sequencing (CITE-seq) technology pose new computational challenges, and there is currently a dearth of rigorous mathematical tools for analyzing the data. This work utilizes concepts and ideas from Riemannian geometry to remove batch effects between samples and develops a statistical framework for distinguishing positive signals from background noise. The strengths of these approaches are demonstrated on two independent CITE-seq data sets in mouse and human. Python source code implementing the algorithms is available at https://github.com/jssong-lab/SAGACITE.

## I. INTRODUCTION

In recent years, single-cell analysis has undergone immense and rapid progress, continuing to transform our understanding of the diversity, development, and cooperation of distinct cell types in various tissues. It is now possible to measure the level of messenger RNAs (mRNAs) in thousands of individual cells via a single experiment of single-cell RNA sequencing (scRNA-seq). Furthermore, multi-omics technologies providing complementary information about the genomic, proteomic, and metabolomic states of single cells are being developed and applied.

Immunophenotyping is the process of classifying immune cells, often relying on the detection of cell-surface proteins. For example, fluorescent activated cell sorting (FACS), a commonly used technique, can be performed before scRNA-seq to provide the immunophenotype information of cells. Two new technologies, cellular indexing of transcriptomes and epitopes by sequencing (CITE-seq) [1] and RNA expression and protein sequencing (REAP-seq) [2], now allow simultaneous performance of immunophenotyping and scRNA-seq transcriptomic profiling in single cells. Both methods are designed to detect proteins on the surface of single cells by adding a panel of DNA-barcoded antibodies on top of the existing high-throughput scRNA-seq techniques. The antibodies bind their corresponding surface proteins, and after cell lysis, the DNA barcodes attached to the antibodies are PCR amplified and sequenced along with the mRNAs. Both CITE-seq and REAP-seq use a unique molecular identifier (UMI)-based protocol, which largely reduces amplification biases. In addition to a count matrix for RNAs, the methods yield a matrix of UMI counts – referred to as the antibody-derived tag (ADT) counts in the CITE-seq literature – derived from sequencing the barcodes attached to the antibodies.

Being less prone to “dropout” effects, the ADT count matrix of surface proteins provides useful information about the immunophenotypes of single cells, while posing new computational challenges in data analysis. Similar to other single-cell techniques, sequencing depth differs from cell to cell; a sound model of ADT count data should take the variation in sequencing depth into account. While UMI-based scRNA-seq data can be modeled with negative binomial (NB) or zero-inflated negative binomial (ZINB) models even for heterogeneous cells [3–5], a direct application of the same approach is not ideal for the count matrix of surface proteins, because a subset of the counts typically comes from nonspecific background binding of antibodies [1]. Fortunately, this type of background noise can be assessed by spiking in control cells from another species that normally do not cross-react with the antibodies. We thus develop a rigorous statistical method for fitting null distributions of ADT counts that are from spiked-in cells, and use it to call positive signals of a protein in the native cells at an adjustable false discovery rate (FDR); to our knowledge, a rigorous statistical framework for such hypothesis testing is not yet available. As model fitting could be adversely affected by systematic differences in measurement between samples, potential systematic biases should be removed prior to model fitting. To accomplish this task, we view single cells as points on a Riemannian manifold and apply ideas from differential geometry to develop a method for removing batch effects between samples.

In this paper, we first introduce the notion of high-dimensional Riemannian manifold endowed with the Fisher-Rao metric, and apply the idea to map the immunophenotype profiles of single cells to a hypersphere. The gist of our batch correction approach relies on the intuition that on this hypersphere, the distribution of points corresponding to spiked-in control cells should be similar between independent samples. We will thus remove sample-specific biases by aligning the center of mass (COM) of spiked-in cells from each sample to a consensus COM. We then apply the same aligning transformation learned from the spike-in data to the native species data, thereby removing potential systematic biases present in both spiked-in and native cells of a given sample. Main computational challenges lie in computing the COM of a point cloud on the hypersphere and “parallel transporting” the point cloud along a specific path connecting the old and new COM, according to some notion of geometry defined on the manifold. Finally, we implement expectation-maximization (EM) algorithms to fit the parameters of our null models describing nonspecific binding of antibodies, and perform statistical tests to detect signals, while keeping the false discovery rate (FDR) under control.

## II. RESULTS

We have applied our geometric and statistical methods to the following two CITE-seq data sets: (1) the public data set of human cord blood mononuclear cells (CBMC) [1], with a low level (~ 5%) of spiked-in mouse control cells; (2) our own data set consisting of 6 samples, each containing immune cells isolated from mouse skin after topical treatment with either inflammation-inducing oxazolone (OXA, 3 mice) or ethyl alcohol (EtOH, 3 mice) as control, as well as a small percentage (∼ 5%) of human HEK293 cells spiked in after the treatment and isolation of the mouse cells [6].

In the first data set, individual cells’ RNA expression (scRNA-seq) profiles were used to identify the mouse spiked-in cells and classify the human cells into distinct cell types (Appendix A); in the second set, scRNA-seq data were used only to identify the human spiked-in cells. For both data sets, the relatively small number of spiked-in cells from a distinct species were identified and separated by calculating the percentage of total RNA counts mapped to the native vs. spiked-in species’ genome (Appendix A). Our approach utilizes the geometric and statistical information contained in spiked-in cells to model the background noise and systematic batch effects in the count data of cell-surface proteins.

Each experiment under consideration has a fixed list of *M* DNA-barcoded antibodies added before sequencing, where each antibody primarily binds its corresponding cell-surface protein, although some non-specific binding manifested as background noise may also be possible. This experimental design determines the list of *M* cellsurface proteins, the abundance of which is to be measured by sequencing the DNA barcodes of corresponding antibodies. For a set of *N* single cells sequenced, a subset of *D* proteins selected from the whole list yields an *N* × *D* matrix of count data,

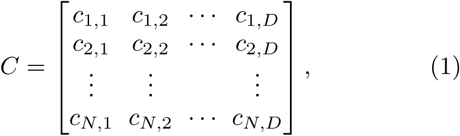

where 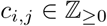 denotes the UMI count, for the *i*-th cell, of the *j*-th protein in the selected subset. A row vector in this matrix thus contains the immunophenotype information of the corresponding cell. In our analysis, the chosen set of *N* cells could be all the cells sequenced in an experiment or only a subset, representing a certain species or a particular inferred cell type. Similarly, the dimension *D* could be equal to the total number *M* of assayed surface proteins, or it could be chosen to be smaller, depending on the biological question of interest.

### A. Mapping immunophenotypes of cells to points on a Riemannian manifold

We first transform the row vector of count data for the *i*-th cell into a probability vector, with the *j*-th component *p*_*j*_ calculated as the fraction 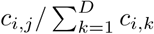. The fraction can be interpreted as the maximum likelihood estimation (MLE) of the probability of finding a certain protein on the *i*-th cell to be the *j*-th protein, given that it is one of the *D* proteins. This transformation maps each cell to a point on the (*D* − 1)-dimensional probability simplex ∆_(*D*−1)_ ⊂ ℝ^*D*^, which, under the coordinate system (*p*_1_,…, *p*_*D*_) of the ambient space, is a polytope satisfying 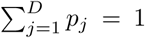 and *p*_*j*_ ≥ 0. On the simplex, the usual Euclidean distance does not properly represent how dissimilar two points are from each other. Hence, we employ mathematical techniques from information geometry and differentiable manifolds to enable the analysis of single-cell data on the probability simplex.

The open probability simplex 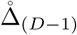, i.e., the relative interior of the probability simplex, satisfying

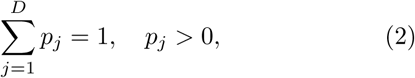

forms a differentiable Riemannian manifold 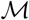 when equipped with the Fisher-Rao information metric [7–9]. A vector 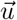 in the tangent space 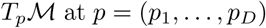 at *p* = (*p*_1_,…, *p*_*D*_) is given by

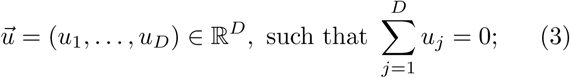

for any 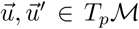, the inner product defined by the Fisher-Rao metric is

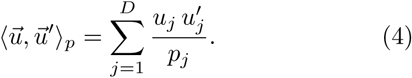

Let 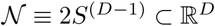 denote a (*D* − 1)-dimensional hypersphere of radius *R* = 2 centered at the origin, and 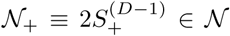 the positive orthant of the hypersphere. It is well known [7–9] that the open probability simplex can be isometrically mapped onto 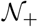 via the diffeomorphism

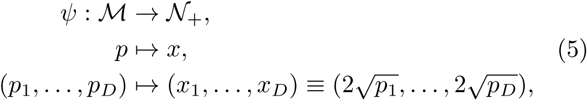

where 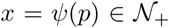 has coordinates (*x*_1_,…, *x*_*D*_) satisfying

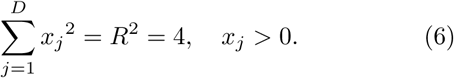

The tangent space at 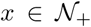 can be obtained as the image of the differential of *ψ*,

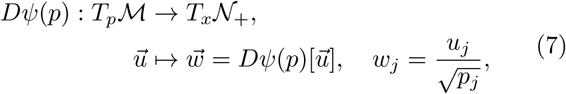

with the standard inner product

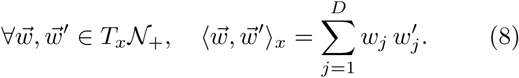

Note that the pullback of this standard inner product on 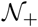 by *ψ* is just the Fisher-Rao inner product on the open probability simplex 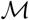.

The entire hypersphere 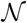 can be regarded as a manifold embedded in the Euclidean space R*D* with the Cartesian coordinates (*x*_1_,…, *x*_*D*_). Unlike the geometry of the open probability simplex, several properties of the hypersphere with the standard induced metric from ℝ*D* facilitate straightforward intuition and calculations. For example, any point 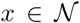 on the hypersphere can be represented as a vector **x** = (*x*_1_,…, *x*_*D*_), such that a vector 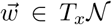 in the tangent space has coordinates **w** = (*w*_1_,…, *w*_*D*_) satisfying 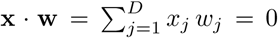, where the dot (⋅) denotes the usual dot product in ∝^*D*^. Furthermore, the geodesic between two points *x* and *y* on a manifold can be derived using the metric-compatible Levi-Civita connection on the manifold. When the manifold is the hypersphere 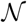 of radius *R* = 2, the geodesic is simply the great arc connecting the two points; that is, with the vector representations **x** = (*x*_1_,…, *x*_*D*_) and **y** = (*y*_1_,…, *y*_*D*_) in the ambient Euclidean space, the geodesic distance between *x* and *y* is given by

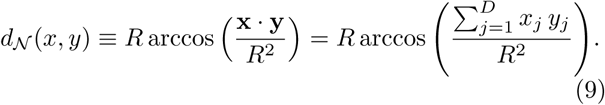

One of the goals in our analysis is to adjust the count data of immunophenotypes by first mapping them to points on the hypersphere 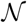, then removing sample-specific biases on 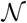 where calculations are simpler, and eventually mapping the corrected points back to count data. As we try to map the count data of cell-surface proteins to the hypersphere, however, there is a small caveat that we need to address. That is, one or more counts of surface proteins might be zero for a cell, and the probability vector will consequently reside on the boundary of the probability simplex, where the Fisher-Rao metric is not defined. Suppose in the probability vector (*p*_1_,…, *p*_*D*_), one component, say *p*_*k*_, is 0. One strategy is to replace it with a small positive number, 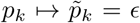, and rescale the remaining components as 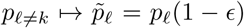, so that the normalization 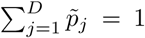 is still preserved. The probability vector now resides on 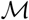 and can be mapped to a point (*x*_1_,…, 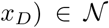 with 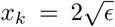. The distance from this point to any other point on 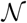 is well defined; taking the limit *ϵ* → 0, the distance remains finite as the point is pushed to the boundary of the positive orthant with its components being *x*_*k*_ = 0 and 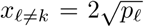. The argument can be generalized to the case where there are more than one protein with zero UMI counts.

In summary, given a selected list of *D* surface proteins and the *N* × *D* count matrix, each row [*c*_*i*,1_,…, *c*_*i,D*_] representing the immunophenotype of the *i*-th cell can be mapped to a point on a (*D* − 1)-dimensional hypersphere of radius 2, with coordinates (*x*_1_,…, *x*_*D*_) given by

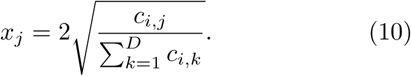

Figure 1 demonstrates two distinct mappings of the indicated cell types from the human CBMC data set. For the sake of visualization, we have chosen the dimensionality to be *D* = 3. For the list {CD3, CD19, CD56}, we observe that most T cells reside in one corner of the positive-orthant hypersphere with a large CD3 component, while most B cells are in another corner with a large CD19 component, both forming densely packed point clouds clearly separated from other cell types and from each other. For the complementary list CD3, CD19, CD56, we see a further separation of CD4+ T cells and CD8+ T cells (although some of them seem to have been misclassified). By contrast, the spiked-in mouse cells and the human erythrocytes (red blood cells) do not possess those human-specific surface proteins expressed on immune cells; therefore, their count data only come from background non-specific binding to the DNA-barcoded antibodies, and their corresponding point clouds lie far away from any of the corners or edges and mostly overlap with each other. In the following sections, we will introduce a method that utilizes the data of spiked-in control cells to remove systematic differences between samples and to model the “noise” of non-specific binding.

**FIG. 1.**
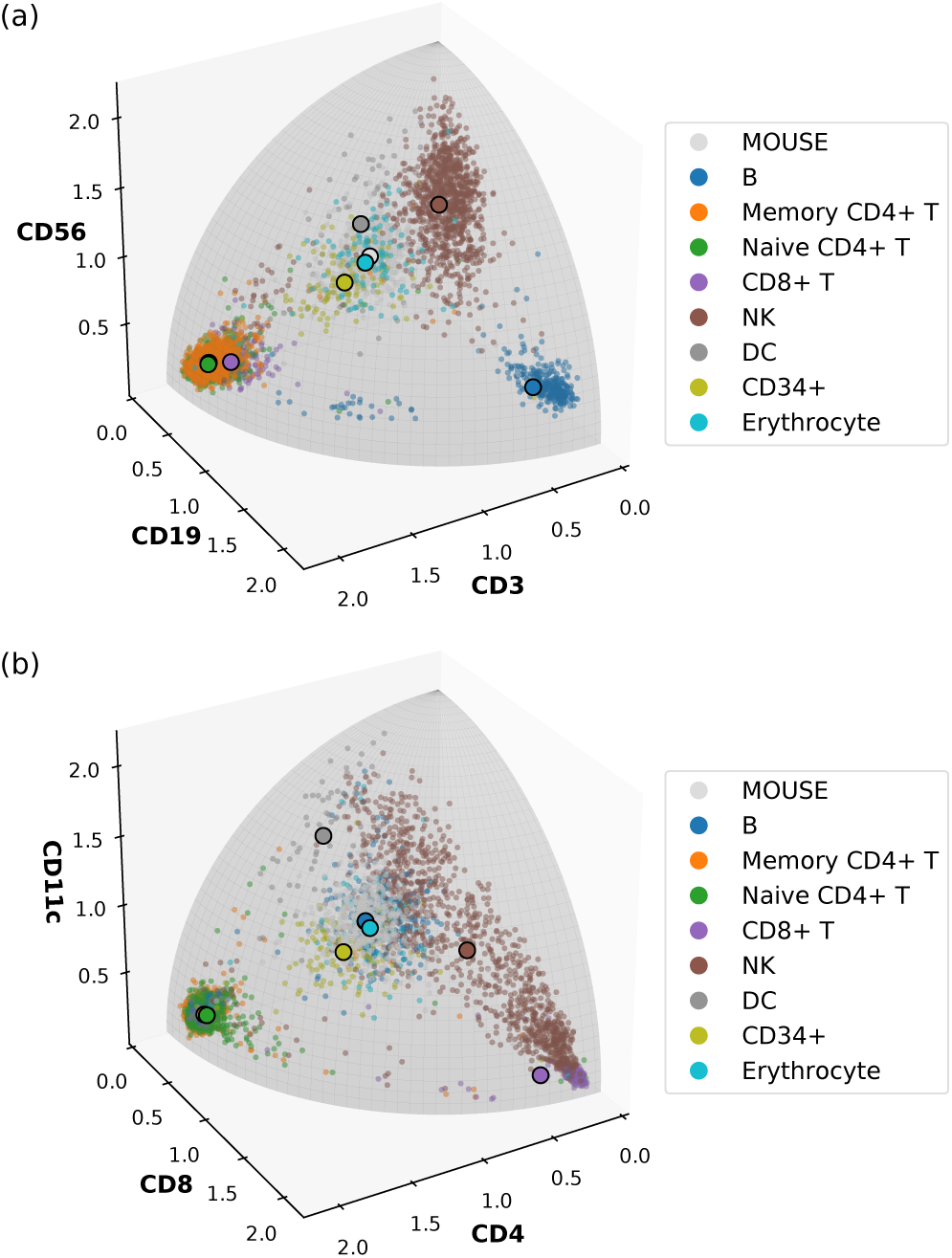
Examples of mapping the surface protein count data of human CBMC with spiked-in mouse cells to a threedimensional sphere of radius 2. (a) The list of selected proteins is {CD3, CD19, CD56}. (b) The list of selected proteins is {CD4, CD8, CD11c}. In both cases, each distinct cell type is displayed with the color indicated in the legend. NK and DC denote natural killer cells and dendritic cells, respectively. Small dots denote individual cells, and large dots with black outlines denote the Riemannian mean of the point cloud of each cell type.

### B. Computing the Riemannian mean and removing batch effects on the hypersphere

With the immunophenotypes of single cells mapped to points on the hypersphere, we can adjust the points from different experiments and remove sample-specific biases by employing the idea, from statistics, of standardizing data distributions by aligning their mean vectors. On a Riemannian manifold, Fréchet mean generalizes the notion of Euclidean mean [10]. It is also often referred to as the Karcher mean, Riemannian center of mass, or Riemannian mean in the literature [9, 11, 12]. In this work, we will simply use the term Riemannian mean and adopt the recent numerical algorithm for computing the Riemannian mean of a set of points on the hypersphere [9].

Computing the COM of a point cloud on a Riemannian manifold involves minimizing an objective function consisting of the pairwise distance between the candidate COM and every point mass in the collection, thus requiring an investigation of the shortest distance between two given points or, equivalently, the geodesic path connecting them. On the (*D* − 1)-dimensional hypersphere 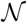 of radius *R*, a geodesic *g* parameterized by *t*, satisfying the conditions 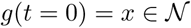 and 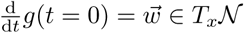, is given by

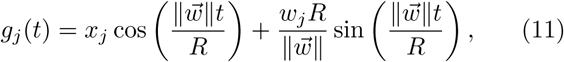

where the embedding coordinates of *x* and *g* in ℝ*D* are **x** = (*x*_1_,…, *x*_*D*_) and **g** = (*g*_1_,…, *g*_*D*_), respectively, and 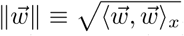. Taking *t* = 1, we obtain the exponential map on the hypersphere, 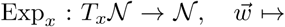 *y* = *g*(1), with the corresponding vector in ℝ^*D*^ given by

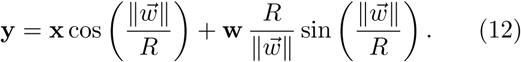

The inverse of the exponential map, 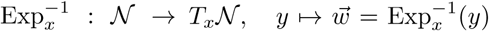, can be easily computed by the Gram-Schmidt process and is given in the embedding Euclidean coordinates by

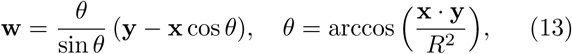

for which *θ* = 0 is reached if and only if *s* = *r*, and in that case, 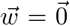 is obtained by taking the limit *θ* → 0. It follows that the norm of the vector 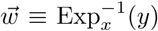 is equal to the geodesic distance between the two points on the hypersphere (9),

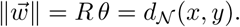

Note that because the exponential map commutes with isometry, the exponential map and the inverse exponential map on the open probability simplex 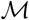 can be obtained from these results on the hypersphere by using *Dψ* and (*Dψ*)−^1^.

We can now define the Riemannian mean on a manifold, generalizing the Euclidean COM as follows: given a set of *N* data points {*y*^(1)^,…, *y*^(*N*)^} with corresponding masses {*m*_1_,…, *m*_*N*_} on the hypersphere, the Riemannian mean 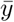 of the collection of point masses is

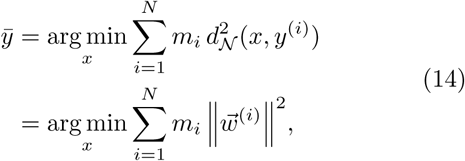

where 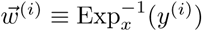 is the inverse exponential map at *x*. The constrained gradient condition with respect to *x* then reads

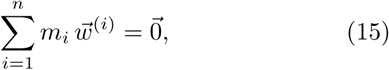

and the Riemannian mean on the hypersphere can be attained numerically in iterative steps until this condition is approximately satisfied [9]. Mapping back the resulting mean to the probability simplex 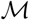 via the inverse isometry *ψ*^−1^ yields the corresponding Riemannian mean on 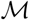. In our biological application, we take all point masses to be 1.

As the human cells in the CBMC data are labeled with cell type information inferred from the transcriptome, we have computed the Riemannian mean for each cell type, as well as the spiked-in mouse cells. The result depends on the choice of surface protein subsets. In Fig. 1, the Riemannian mean of each cell type is shown as a large dot with black outline. We see that the Riemannian mean is a good representative of a densely packed point cloud on the sphere. Figure 2 shows the components of the Riemannian mean for *D* = *M* (all the proteins) and *D* = *M* − 1 (CD45 excluded). We see that, in this case, excluding CD45 increases the contrast of specific markers associated with each cell type, due to the fact that CD45 is generally high in human immune cells and may suppress the resolution of other cell-surface markers specific to certain immune cell subtypes. We also see that CD10, CCR5, and CCR7 are not biological markers for any of the cell types, consistent with the result in [1]. This analysis illustrates how the Riemannian mean summarizes a set of homogeneous cells, and this idea will guide our method for removing sample-specific biases.

**FIG. 2.**
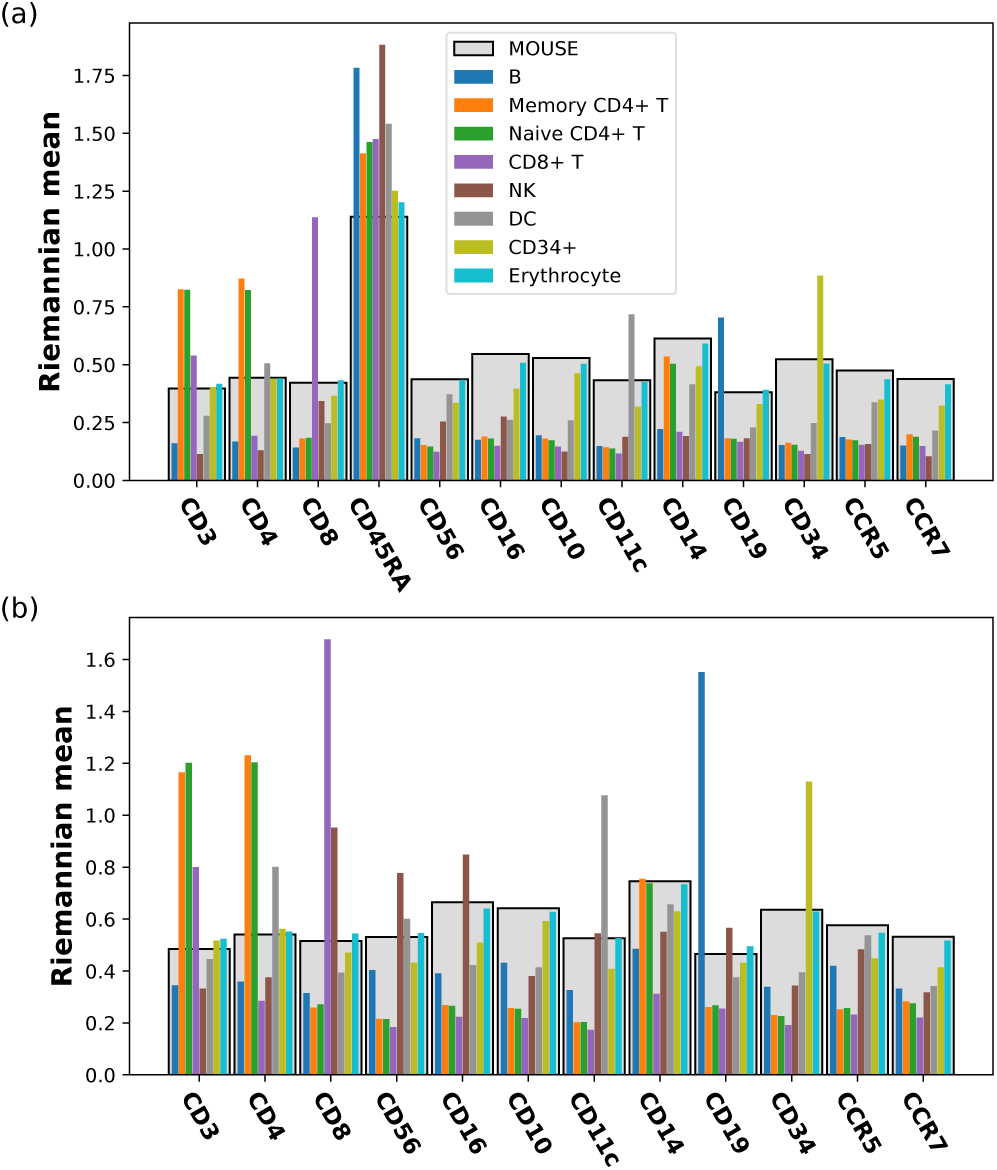
Riemannian mean calculated from the surface protein count data of each indicated cell type in the human CBMC data set. (a) All proteins are included (*D* = 13). (b) CD45RA is excluded (*D* = 12). In both ways of mapping, the components of the Riemannian mean correspond to the height of the bars; the light gray bars in the back represent the spiked-in mouse data, while the thin bars in the front represent the different cell types in human blood, with their order and colors indicated in the legend.

For our own data set of immune cells isolated from the mouse skin, plus spiked-in human cells, we have only identified the species without further classification into distinct cell types. Figure 3 shows the components of Riemannian mean for the mouse and human cells in each of the 6 samples. We observe not only biological differences caused by the different treatment conditions – e.g., the enrichment of CD11b in OXA-treated mouse cells – but also some systematic differences between samples subjected to the same treatment – e.g., CD69 being much higher in EtOH2 than in EtOH1 and EtOH3 for both mouse and human data. For the spiked-in human cells, all the count data should in principle come from non-specific binding, but the Riemannian mean of some samples are very much separated from the rest. In fact, certain surface proteins (e.g., CD69, CD44, CD134, and CD86) show notable, reproducible skews in both mouse and human cells of the same sample. The pattern of certain surface protein enrichment in spiked-in human cells of a specific sample and the persistence of these biases in the mouse cells of the same sample suggest that there might exist systematic differences between the samples. These differences between samples can be further visualized in the principal component analysis (PCA), where the point clouds of EtOH2, OXA1, and OXA2 are seen not to overlap with the other similarly treated samples for both human and mouse cells [Fig. 4(a,b)]. Systematic differences between samples, also known as batch effects, will prevent comparison between different experiments. We now describe our method for correcting such batch effects by aligning the Riemannian mean of samples and parallel transporting the collection of data from each sample along a geodesic path connecting the old COM and the new aligned COM.

**FIG. 3.**
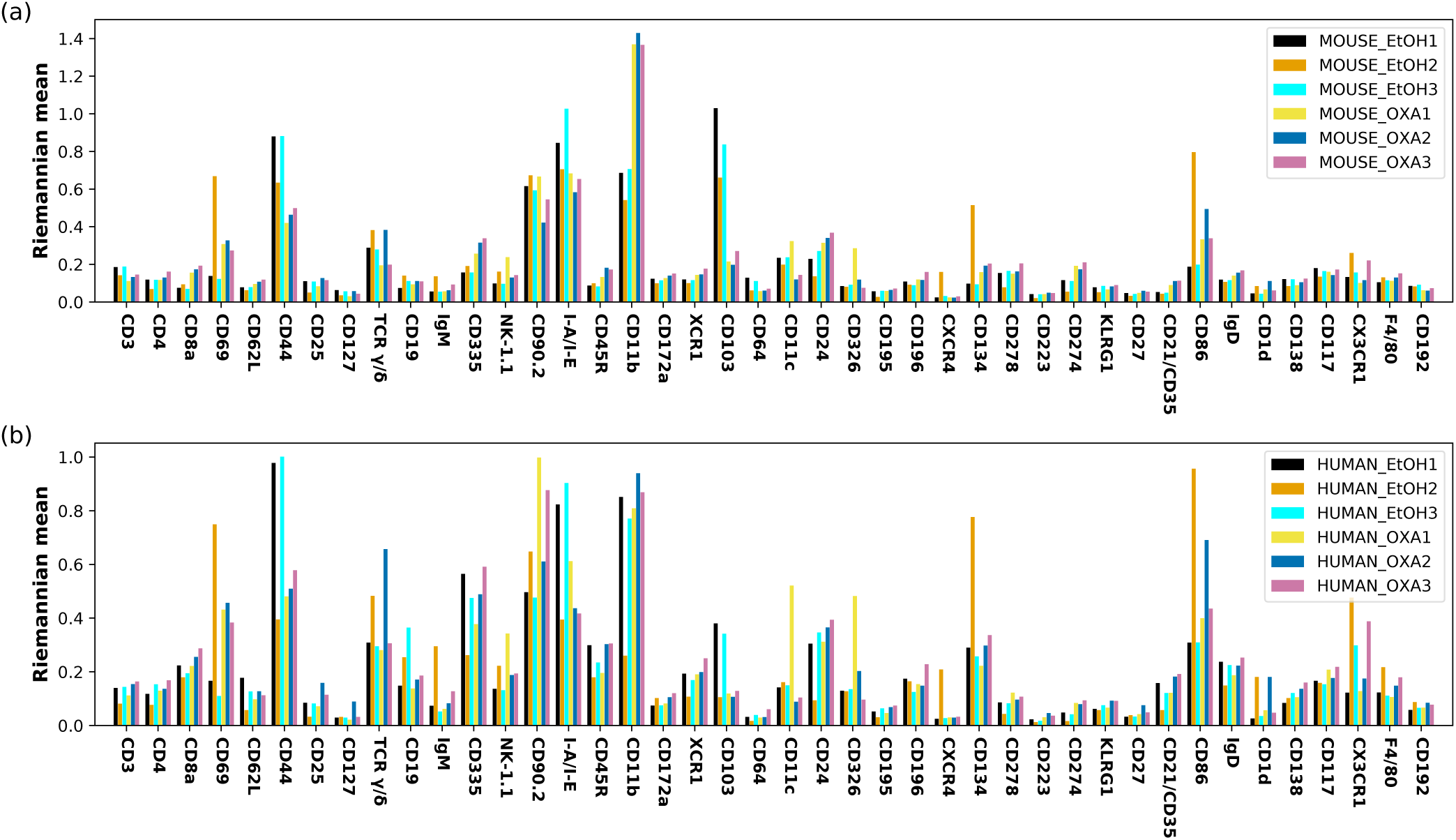
Batch effects within the three oxazolone-treated samples (OXA1,2,3) and the three control samples (EtOH1,2,3) of mouse skin cells. (a) Riemannian mean of the native mouse cells from each sample. (b) Riemannian mean of the spiked-in human cells from each sample. The bar height corresponds to the component of the Riemannian mean in the direction indicated on the *x*-axis.

**FIG. 4.**
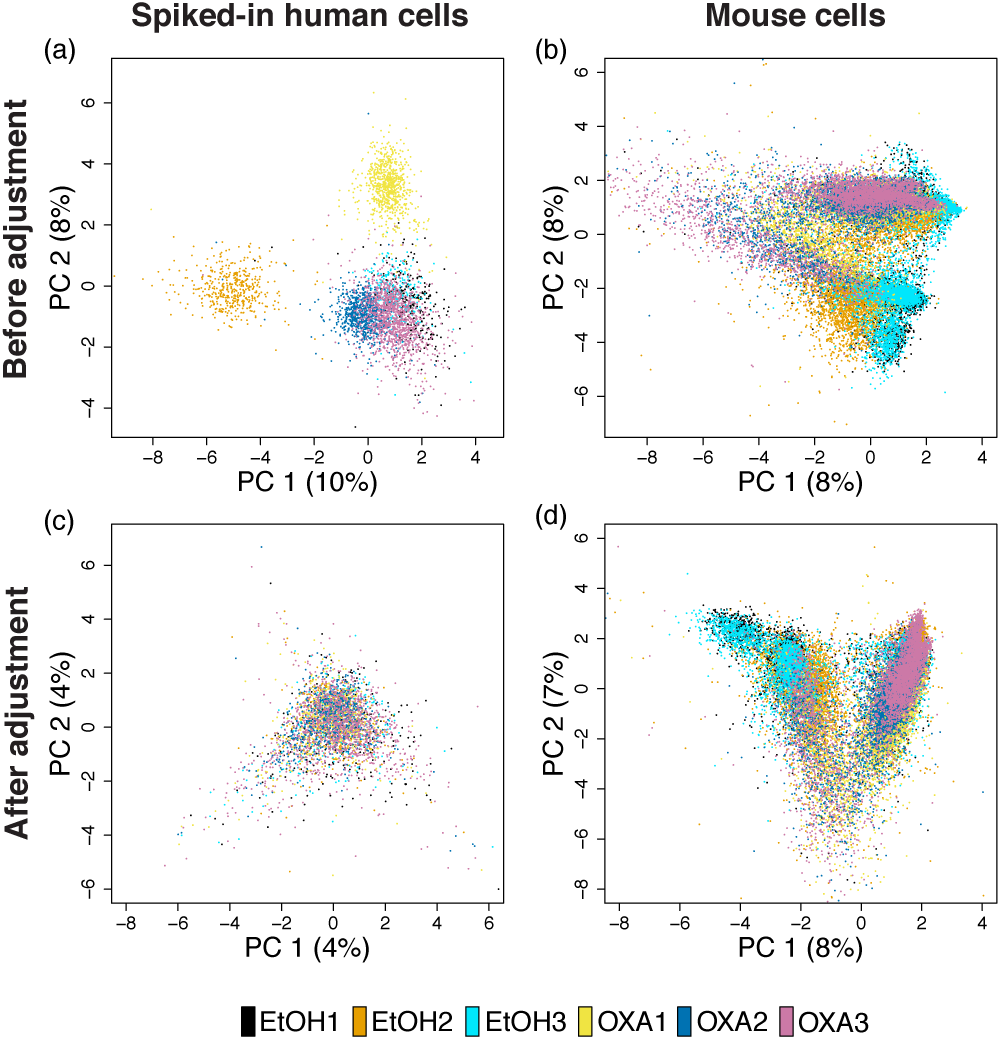
Principal component analysis (PCA) of spiked-in human cells and native mouse cells on the probability simplex before and after batch correction. The single-cell ADT count data of surface proteins were transformed to probability vectors and then projected to the plane spanned by the first two principal components (PCs). (a) Spiked-in human data before batch correction. One control sample (EtOH2) and two treated samples (OXA1,2) are seen to be outliers from the rest. (b) Mouse data before batch correction. The biases observed in (a) are seen to be carried over here. (c) Spiked-in human data after batch correction. Point clouds of all six samples are seen to overlap well. (d) Mouse data after batch correction. Points from the six samples are seen to align well with respect to the two treatment conditions. The variance explained by the first two PCs is also slightly reduced.

In the above discussion of Riemannian mean, we have seen that given a reference point *x*, points *y*^(1)^,…, *y*^(*N*)^ in its neighborhood can be mapped to vectors 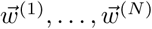 in the tangent space 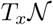 via 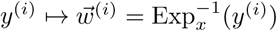, and vice versa. Using the Levi-Civita connection, we propose to parallel transport these vectors along the geodesic path from *x* to a new reference point *z*, and then retrieve points in the neighborhood of *z* via the exponential map acting on the transported vectors lying in 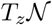. On the hypersphere embedded in ℝ*D*, this transformation is equivalent to a rotation in the plane spanned by the vectors **x**, **z** ℝ^*D*^. Using (12) and (13), we have for the rotation

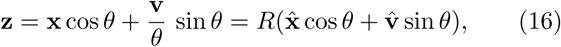

with 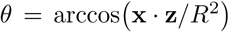, 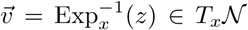, and the unit vectors 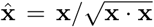 and 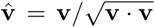. For a point *y*^(*i*)^ in the neighborhood of *x*, the corresponding point transported to the neighborhood of *z* is given by

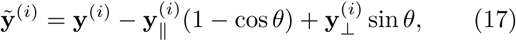

in which

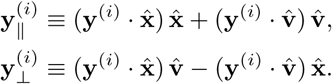

This approach thus enables a method of correcting batch effects by aligning the Riemannian mean *x* of a point cloud from each sample to a consensus reference point *z* and thereby transporting each point cloud to the neighborhood of *z*. As previously discussed, the spiked-in human cells in each mouse sample are supposed to measure “noise” from non-specific background binding and should be similarly distributed on the hypersphere. We have chosen the consensus reference point *z* to be the Riemannian mean of the aggregated human cells from three samples for which the point clouds of human cells mostly overlap, namely EtOH1, EtOH3, and OXA3 [Fig. 4(a)]. The points transported to this reference point *z* represent the immunophenotypes of cells after the batch correction. The same rotation on the hypersphere is now applied to the mouse cells to remove systematic biases between samples.

As an approximate inverse of (10), we calculate the corrected fractions of surface proteins from the corrected coordinates on the hypersphere, and restore the count data by multiplying the fractions by the total number of UMI counts and rounding the results to nearest integers. It is possible that after the batch correction, some points on the hypersphere could have small negative coordinate components; i.e., some points may be moved out of the positive orthant after the rotation. As the components of a probability vector can never be negative, we force the negative components in (*x*_1_,…, *x*_*D*_) to be zero, and rescale other components so that the probability normalization still holds. In our experience, this thresholding has a negligible effect on the downstream analysis. We see in Fig. 4(c) and (d) that our batch correction method has successfully removed the differences between spiked-in human cells, and also aligned the mouse samples according to the treatment conditions. We showcase the successful removal of batch effects using four surface proteins, CD69, TCR *γ*/*δ*, CD90.2 and I-A/I-E, in Fig. 5. Before the correction is applied, it can be seen that biases found in the distribution of the spiked-in human cells in an outlier sample is often replicated in that of the mouse cells in the same sample – e.g., CD69 in EtOH2, TCR *γ*/*δ* in EtOH2 and OXA2, CD90.2 in OXA1, and I-A/I-E in EtOH3 and OXA1. These systematic biases are largely removed in both human and mouse cells by our batch correction method.

**FIG. 5.**
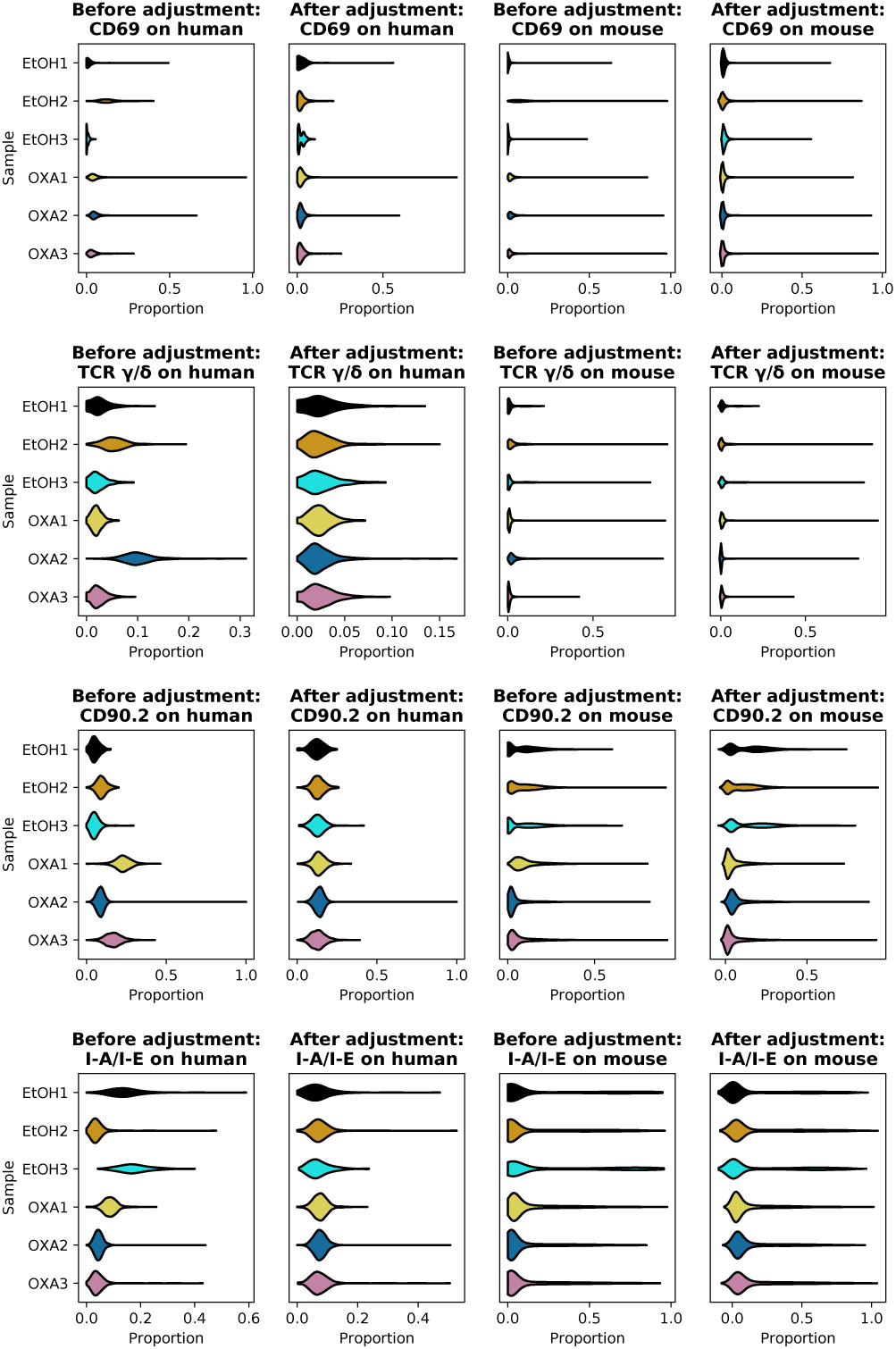
Effects of batch correction on the six samples of mouse skin cells with spiked-in human cells. The data points on the hypersphere either before or after the batch correction are mapped back to the probability simplex. Distributions of proportion for human and mouse cells in each of the six samples are shown for the four selected surface proteins CD69, TCR *γ*/*δ*, CD90.2, and I-A/I-E.

### C. Fitting the null model and performing statistical tests on count data

We now present a statistical framework for testing the significance of enrichment of a specific surface protein in the sequencing of a native cell, compared to the null distribution of read counts for that protein in the population of spiked-in cells. For the *j*-th surface protein, we build the null model on the *j*-th column, {*c*_1,*j*_, *c*_2,*j*_,…, *c*_*N,j*_}, of the count matrix for spiked-in cells. In the following, we will focus on one surface protein at a time and omit the subscript *j* indexing surface proteins to simplify notation.

When the number of zero counts is small for the surface protein under consideration, we propose to model the null distribution by a generalized form of negative binomial (NB) model, with cell-specific relative size factors {*t*_1_, *t*_2_,…, *t*_*N*_} capturing differences in individual cells’ sequencing depths. The idea is similar to that described in [4] for analyzing scRNA-seq UMI counts, but instead of regressing a generalized linear model (GLM), we will utilize the expectation-maximization (EM) algorithm to estimate the parameters in the NB model [13]; we will subsequently show that the algorithm can also be modified to estimate the paramters of zero-inflated models that are suitable for sparse data. Once the model parameters are determined, we will then use the null model to perform statistical tests on the count data of surface proteins in native cells, distinguishing “signal” from “noise” while keeping the false discovery rate (FDR) under control.

For the *i*-th cell, we denote the surface protein count random variable as *y*_*i*_ and its actual observed value as 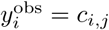. (This notation is not to be confused with the previous section’s Cartesian coordinates **y** = (*y*_1_,…, *y*_*D*_) on the hypersphere.) The relative size factor *t*_*i*_ is a measure of the cell’s sequencing depth covering all surface proteins, relative to a typical sequencing depth among all *N* cells. The calculation of *t*_*i*_ is discussed in Appendix D, where we offer two choices of definition, (D6) and (D7). We use the expression of *t*_*i*_ defined in (D7) in this paper, unless stated otherwise.

The NB probability distribution for a count random variable *y*_*i*_ with a predetermined relative size factor *t*_*i*_ is

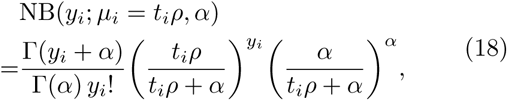

parametrized by a cell-specific mean µ_*i*_ and a universal ‘stopping-time’ parameter *α*, with the mean being E[*y* ] = *µ* = *t*_*i*_ *ρ*, and the variance 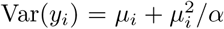. For our EM implementation, it is instructive to view the NB distribution as an infinite mixture of Poisson distributions with mixing coefficients given by the Gamma distribution. This paper uses the following conventions:

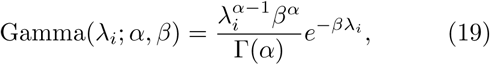

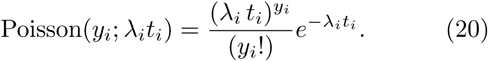

The shape parameter *α* and rate parameter *β* for the Gamma distribution are the same for all *N* cells in the set. In the mixture model, once λ_*i*_ is sampled from the Gamma distribution for the *i*-th cell, the mean of the Poisson distribution is determined by the product of λ_*i*_ and the cell-specific size factor *t*_*i*_, independent of the parameters (*α*, *β*); we get the NB probability distribution under a reparametrization *ρ* = *α*/*β*, as

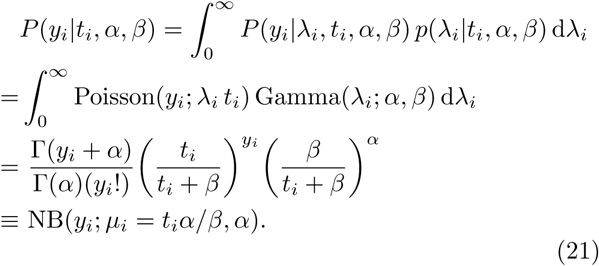

In this formulation, the NB probability distribution arises by marginalizing the hidden variable λ_*i*_ from the joint distribution of (*y*_*i*_, λ_*i*_). In this paper, we use *P* to denote both probability mass functions of discrete random variables and joint distributions of discrete and continuous random variables. To apply the EM algorithm to infer *α* and *β*, we need not only the joint distribution

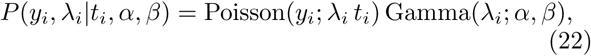

but also the posterior density of λ_*i*_, computed by applying the Bayes’ rule as

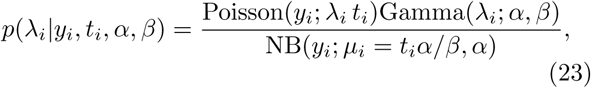

which is also Gamma distributed. Once the maximum likelihood estimation (MLE) of the parameters (*α*, *β*) is attained, we can interpret the mean of this posterior distribution as the expected surface protein levels of single cells.

For the set of *N* homogeneous spiked-in cells, with count variables **y** = {*y*_1_, *y*_2_,…, *y*_*N*_}, their observed values being 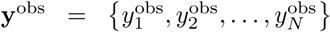, and corresponding size-factors *t* = {*t*_1_,*t*_2_, …, *t*_*N*_}, we have 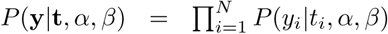. Denoting **λ** = {λ_1_, λ_2_,…, λ_*N*_} to be the set of hidden variables for the *N* cells, the full joint distribution is 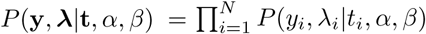, and we have

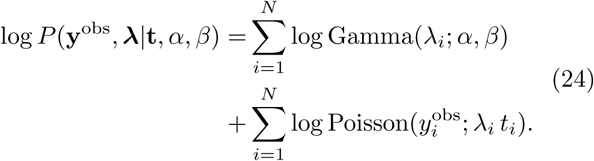

By taking expectation of (24) with respect to the posterior distribution of **λ** given in (23), we then apply an implementation of the EM algorithm to obtain the maximum likelihood estimates of the model parameters *α* and *β* (see Appendix B for details) [13].

When an experiment produces 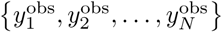 containing a relatively large number of zero counts, we use a zero-inflated negative binomial (ZINB) model, defined by

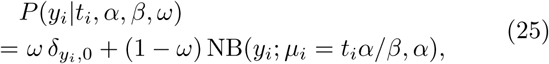

where *δ*_*y*,0_ = 1 for *y* = 0, and 0 otherwise. The new parameter *ω* is the probability of a “dropout” event in the measurement. Upon some modification, the above EMalgorithm can be used to obtain the maximum likelihood estimates of *α*, *β*, and *ω* (see Appendix C).

Note that the NB distribution is a special case of the ZINB distribution with *ω* = 0. We have observed that if the NB model is sufficient for modeling the observed counts, then fitting a ZINB model will give *ω* → 0. However, fitting a ZINB model will take longer time, and one may wish to choose a particular model based on the data at hand. The human CBMC dataset is not sparse, with only a few zero counts; we have thus chosen to fit a NB model for each of the *M* = 13 surface proteins. By contrast, the count data of mouse skin cells are sparse, with the rates of zeros sometimes as high as 70%, and we have chosen to fit a ZINB model for each of the *M* = 42 proteins.

Fitting the NB or ZINB distribution on the count data of each surface protein from spiked-in cells yields a null model, from which we can compute the *p*-values for the observed counts in native cells and thus distinguish potential “signals” from “noise” in native cells. In Fig. 6(a), we show the null model fitted on the count data of human CD3 observed on the spiked-in mouse cells, with the distribution of *p*-values being nearly uniform and thus indicating a reasonable fit. The *p*-values of the human cells show a clear distinction from the null model and show an enrichment of cells having a significant signal of CD3 (small *p*-values). Comparing the count data from native cells to the null distribution of spiked-in cells introduces a problem of multiple hypothesis testing. To control for the false discovery rate (FDR), we use the Benjamini–Hochberg (BH) procedure [14], as implemented in the r package ‘stats’ [15]. The adjusted *p*-values for mouse and human data are shown in Fig. 6(b). We would decide a cell as having a significant signal for a surface protein when the corresponding adjusted *p*-value is smaller than the chosen FDR threshold.

**FIG. 6.**
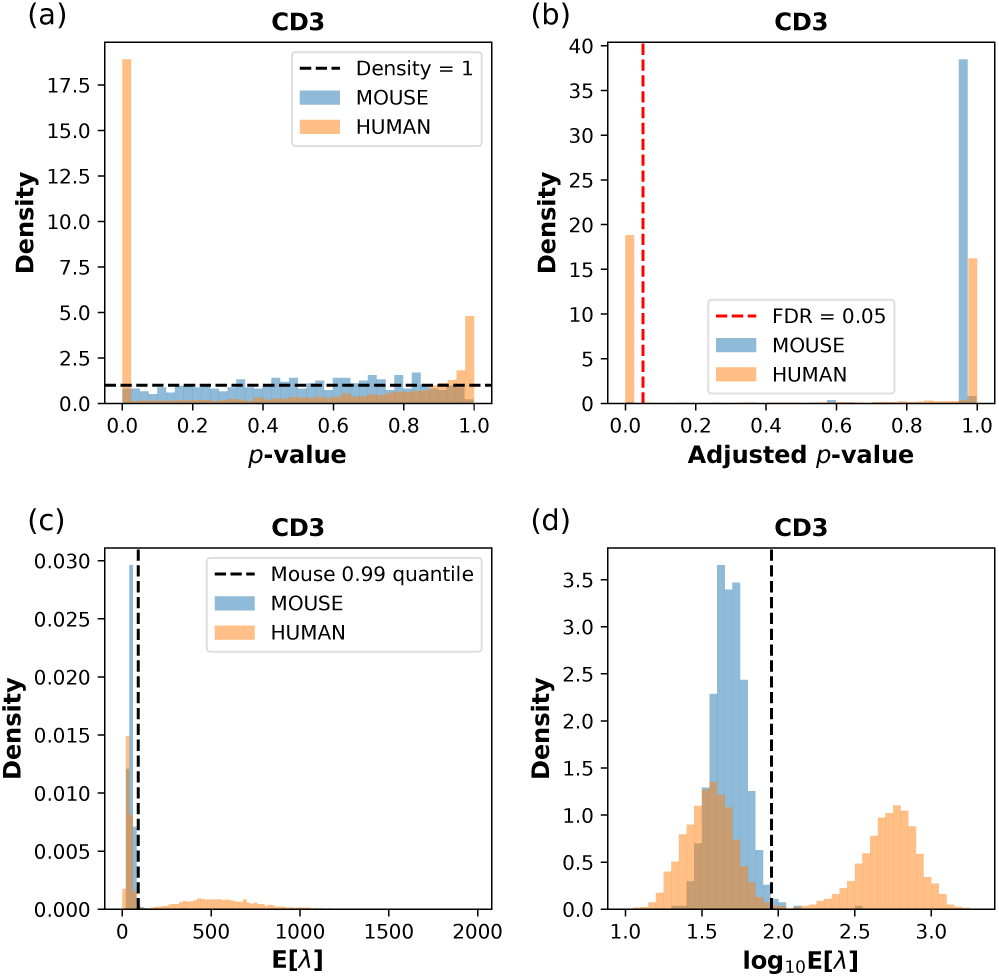
Fitting the NB model on spiked-in mouse cells in the CBMC data set, and performing statistical tests and data transformation with the estimated model parameters. The surface protein is chosen to be (human) CD3, with the fitted model paramters *α* = 10.30, *β* = 0.2074 estimated from the mouse data, and *ω* = 0 fixed for the NB model. (a) The distribution of *p*-values for mouse and human cells calculated from the fitted model. The horizontal dashed line indicates a uniform distribution with constant density 1. (b) The distribution of adjusted *p*-values. The vertical red dashed line indicates the FDR threshold of 0.05; cells to the left of this line are considered as CD3+, and they are all human cells. (c) The distribution of the posterior mean E[λ] for mouse and human cells calculated from the model parameters. (d) The distribution of log10 E[λ] for mouse and human cells. In (c) and (d), the vertical dashed line indicates the 0.99 quantile of the spiked-in mouse data.

The model fitting also provides a method of transforming the count data in a way compatible with the statistical model. The transformed data can be used for downstream analysis, such as correlation analysis, dimension reduction, and unsupervised clustering. With the ZINB parameters (*α*, *β*, *ω*) estimated from a given set of (**y**^obs^, **t**), we can transform a pair (*y′*, *t′*) of observed UMI count and size factor, either used in the model fitting or previously unseen, as

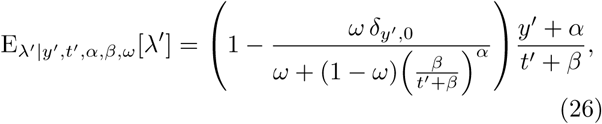

which is roughly the posterior expected Poisson mean of the UMI-count random variable in the ZINB model. For the NB model, we simply need to take *ω* = 0, and the expectation value reduces to (*y′* + *α*)/(*t′* + *β*). The transformed CD3 count data in the CBMC data set are shown in Fig. 6(c) and (d), where the human data are clearly seen to have a mode with high E[λ] signals, well separated from the background distribution of spiked-in mouse cells. Taking the logarithm of E[λ] results in a better visualization, as seen in Fig. 6(d), clearly capturing the bimodal distribution of the CD3 expression level on the surface of human blood cells. For CD3, the number of human cells above the 0.99 quantile of the mouse control distribution approximately coincides with the number of human cells passing the statistical test at the adjusted *p*-value threshold of 0.05 in Fig. 6(b).

## III. DISCUSSION

Inspired by the techniques of differential geometry and stochastic processes often used to model physical systems, we proposed a series of methods for analyzing the count data of surface proteins from CITE-seq. Mapping the count data to a Riemannian manifold, we used the Riemannian center of mass to find an exemplar point that best represents each set of homogeneous control cells on the manifold. We then removed potential batch effects between multiple samples by aligning their center of mass on the Riemannian manifold and built a null model in order to separate significant signals in the count data from the noise of non-specific antibody binding.

To date, CITE-seq analysis lacked a rigorous statistical framework for testing the significance of ADT counts and adjusting for multiple hypothesis testing. Our probabilistic modeling of ADT sequencing addresses this gap and also provides an appealing data representation based on the posterior mean E[λ], which can be used for down-stream analyses such as clustering and visualization. Inheriting the parameters from the (ZI)NB model fitting makes the transformation easily interpretable and compatible with the proposed statistical hypothesis testing framework. Further details and comparison with other data transformation (normalization) methods are discussed in Appendix D.

Unlike the original approach [1], some CITE-seq data may lack a spike-in control from another species. In those cases, we recommend first finding a set of non-immune cells (e.g., erythrocytes in the blood, and keratinocytes in the skin [16, 17]) that are transcriptomically distinct from the rest of the cells, and then using the set to build the null model for immunophenotype profiling. If this strategy is not feasible, then an unsupervised method could be developed to distinguish signal from noise by fitting bimodal or multimodal distributions.

As previously mentioned, parallel transporting the immunophenotypes of cells on the hypersphere might move some cells slightly out of the positive orthant. We here addressed the issue by setting the small negative components to zero and rescaling the rest of the components to preserve the normalization condition. Even though this simple correction method did not noticeably affect the neighborhood structure of the point clouds in our data, future studies would be needed to develop a more rigorous geometric construction that can handle these cells.

Our method of batch correction, upon some modification, may be also applicable to other types of count data; e.g., other multi-omics count data that complement the scRNA-seq assay, and count data used for topic modeling in text mining. A potentially interesting direction for future investigation would be integrating the geometric and statistical methods directly on a Riemannian data manifold.

## ACKNOWLEDGMENTS

This project was supported in part by funds from NIH K08AR067243 (JBC), NIH R01CA163336 (JSS) and the Grainger Engineering Breakthroughs Initiative (JSS).

## Appendix A: Data set preparation

The CITE-seq data set of human CBMC and spiked-in mouse cells was obtained from the Gene Expression Omnibus under the accession number GSE100866. For the scRNA-seq data, we followed the suggested procedures of normalization, feature selection, dimensional reduction, and Louvain clustering in Seurat v3 [18]. The cell labels were determined from the list of biomarkers detected for each cluster using Seurat, as in [1]. Furthermore, we summed up the RNA counts mapping to the mouse genome, and calculated the percentage of mouse gene counts with respect to the total RNA counts; the putative single cells with a percentage of mouse genes from 5% to 95% were filtered out, as they might be doublets of cells from the two species. The cells with larger than 95% mouse genes were labeled as mouse cells. A tSNE plot of the transcriptomic data with labeled cells is shown in Fig. 7(a). For a clear demonstration of our analysis, we have chosen human cells with labels only from the following eight cell types: B cells, memory CD4+ T cells, naive CD4+ T cells, CD8+ T cells, natural killer (NK) cells, dendritic cells (DCs), CD34+ cells and erythrocytes. The cells labeled as CD14+ monocytes, CD16+ monocytes, megakaryocytes, plasmacytoid dendritic cells (pDCs), and multiplets were all filtered out. The full list of 13 cluster of differentiation (CD) proteins measured in the experiment is *{*CD3, CD4, CD45RA, CD56, CD16, CD10, CD11c, CD14, CD19, CD34, CCR5 (CD195), CCR7 (CD197), all of which are shown on the *x*-axis in Fig. 2.

**FIG. 7.**
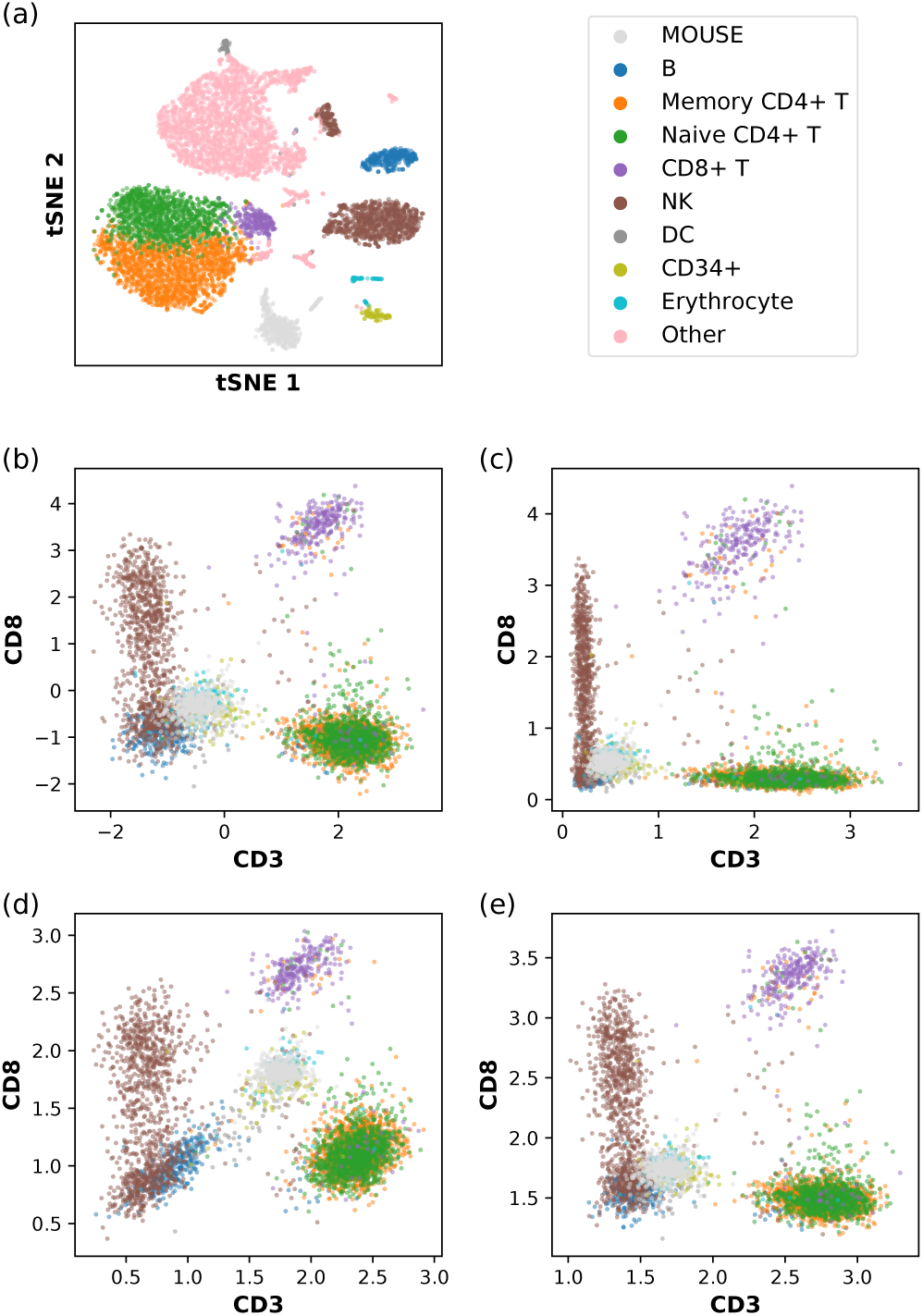
Data transformation applied to human CBMC with spiked-in mouse cells. (a) tSNE plot of the single-cell transcriptomic (scRNA-seq) data. The RNA count data have been log-normalized, as described in Eq. (D1), and compressed using a dimensional reduction method (Appendix A). The indicated color scheme for cell types is carried over to (b,c,d,e). NK and DC denote natural killer cells and dendritic cells, respectively. CD14+ monocytes, CD16+ monocytes, megakaryocytes, and plasmacytoid dendritic cells (pDCs) are grouped into the category “Other” and omitted in other panels. (b) A version of the centered log ratio (CLR) transformation of the single-cell immunophenotype data, as described in Eq. (D2). (c) Another version of the CLR transformation of the single-cell immunophenotype data, as described in Eq. (D3). (d) Our data transformation method using the relative size factor *t*_*i*_ = *a*_*i*_/*a*_0_, with *a*_*i*_ being the arithmetic mean of count per protein, as described in Eq. (D6). (e) Our data transformation method using the relative size factor *t*_*i*_ = *g*_*i*_/*g*_0_, with *g*_*i*_ being the geometric mean of count (plus one pseudocount) per protein, as described in Eq. (D7).

Processed tables of ADT counts in murine skin cells and spiked-in human embryonic kidney 293 (HEK293) cells [6] are available at https://github.com/jssong-lab/SAGACITE. The samples OXA1, 2, and 3 were from the ear skin of three different mice treated with inflammation-inducing oxazolone, while EtOH1, 2, and 3 were from the ear skin of three different mice treated with ethyl alcohol as control. The immune cells in each skin sample were isolated after enzymatic digestion and then cell sorting using flow cytometry. The HEK293 cells were then spiked in, just before CITE-seq was performed. For each cell, we calculated the percentage of RNA counts mapping to the mouse genome with respect to the total amount of RNA counts; a cell with the percentage greater than 99% was classified as a mouse cell, and a cell with the percentage smaller than 5% was classified as a human cell. No further sub-classification of mouse cells based on the transcriptome was performed.

## Appendix B: EM algorithm for NB model fitting

The posterior density *p*(λ_*i*_|*y*_*i*_, *t*_*i*_, *α*, *β*) of λ_*i*_ defined in (23) satisfies

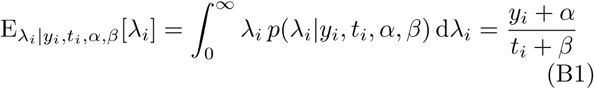

and

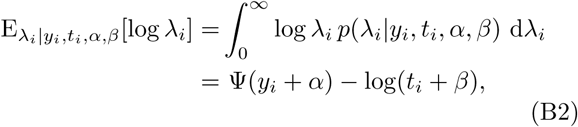

where 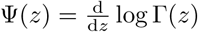 is the digamma function.

For a dataset (**y**, **t**) of *N* independent samples, we define the following sample averages of expectation values:

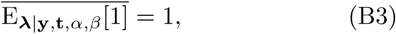

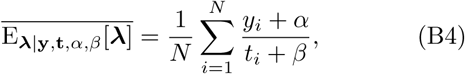

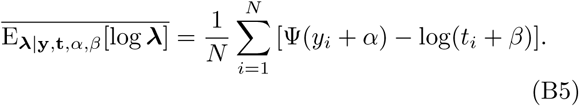

Given the current estimate of (α*, β*), we need to update them as

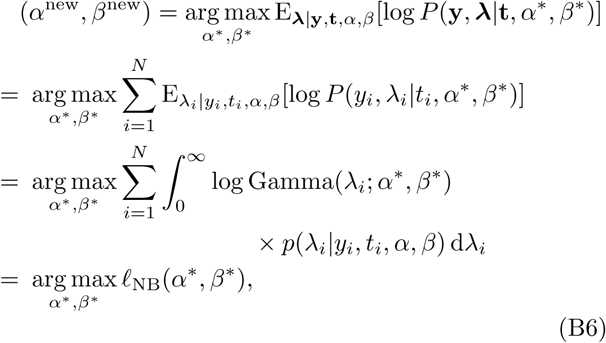

where

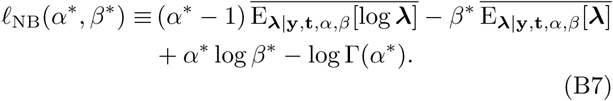

Solving 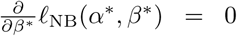 for *β*∗, we get 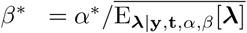. Substituting this expression of *β*∗ into *£*_NB_(*α*∗, *β*∗), we define a new function depending only on *α*∗:

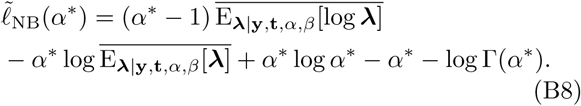

Maximizing this function, we finally obtain the updates

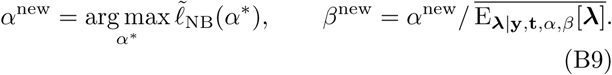

In each iterative step, we compute the optimization of *α*^new^ numerically using a generalized version of Newton’s method which enables faster convergence [13, 19].

## Appendix C: EM algorithm for ZINB model fitting

For the ZINB model, the joint probability function is

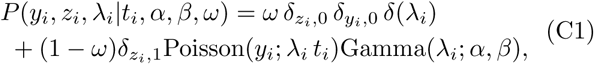

where *z*_*i*_ ∈ {0, 1} is a latent Bernoulli random variable modeling the “dropout” event (*z*_*i*_ = 0). The posterior probability function is

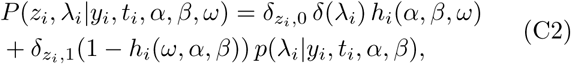

where the posterior density *p*(λ_*i*_|*y*_*i*_, *t*_*i*_, *α*, *β*), for *z*_*i*_ = 1, is defined in (23), and

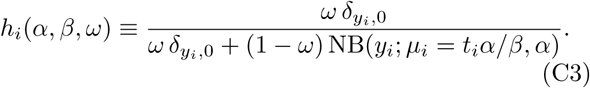

Given the current estimate of (*α*, *β*, *ω*), we need to update them as

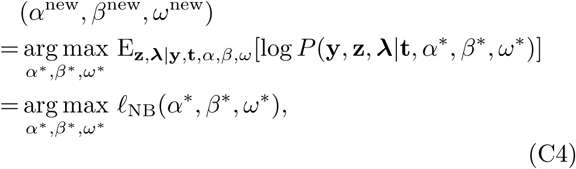

where *£*_ZINB_(*α*∗, *β*∗, *ω*∗), collecting only the terms involving *α*∗, *β*∗, *ω*∗, is given by

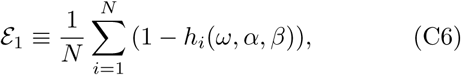

In the last line, **𝓔**_1_, **𝓔**_**λ**_, and **𝓔**_log **λ**_ are defined as follows, similar to (B3), (B4), and (B5):

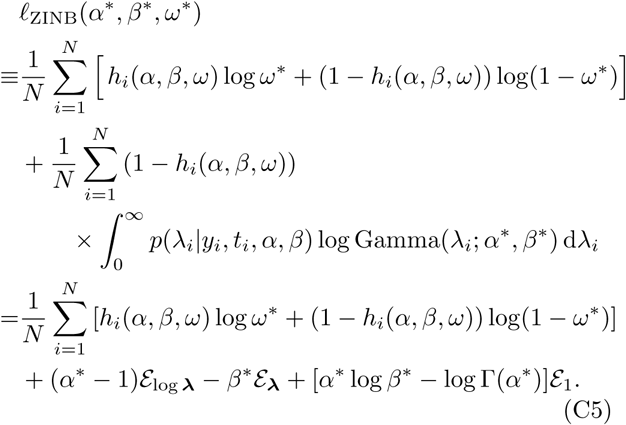

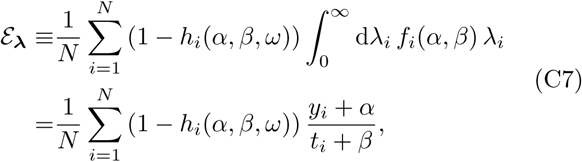

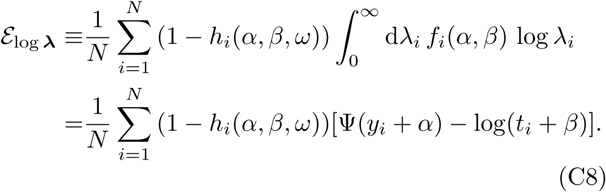

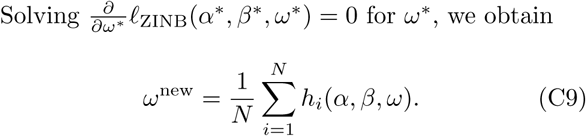

From 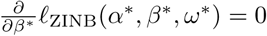, we have *α*\*/*β*\* = **𝓔**_**λ**_/**𝓔**_**1**_. Keeping only those terms involving *α*∗ and *β*∗ in (C5) and substituting *β*∗ = *α*∗ **𝓔**_1_/**𝓔**_**λ**_, we can define a function that depends only on *α*∗ as follows

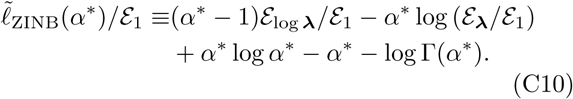

The update now reads

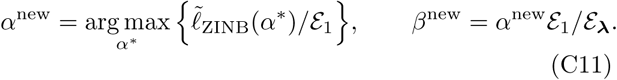

With the ratios **𝓔**_logλ_/**𝓔**_1_ and **𝓔**_logλ_/**𝓔**_1_ calculated using (C6), (C7), and (C8), the resulting optimization is the same as in (B8).

## Appendix D: Comparison of data transformation

The log normalization with a fixed scale factor *s*_0_, commonly used to process scRNA-seq data, transforms the count data (*c*_*i*,1_,…, *c*_*i,D*_) within the *i*-th cell as

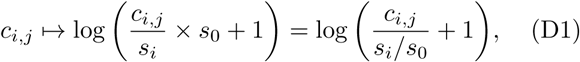

where 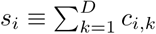 is the total sequencing depth in the *i*-cell, and *s*_0_ can be chosen to be a typical value of *s*_*i*_. Some common choices are *s*_0_ = 1000, 10000, or 100000, depending on the data. We also find it reasonable to choose *s*_0_ as either the arithmetic or the geometric mean of *s*_*i*’_s.

The centered log ratio (CLR) is a related transformation method that is previously used to process the CITEseq count data [1], and is defined as

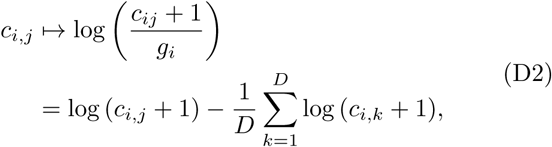

where 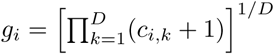 is the geometric mean of the *D* surface protein counts, each adjusted by one pseudocount. It can be interpreted as row mean-centering the table of pseudocount-adjusted log counts, log (*c*_*i,j*_ + 1). Another version of the CLR transformation, implemented in Seurat v3 [18], is given by

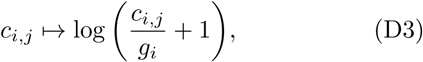

which is not row-centered. Unlike the expression defined in (D2), the alternative form given in (D3) always yields nonnegative values.

We here propose a new data transformation method using the posterior E[λ_*i,j*_] computed using the MLE of ZINB model parameters (*α*_*j*_, *β*_*j*_, *ω*_*j*_) for the *j*-th protein, as given in (26); we take the logarithm for better visualization of the data, the effect of which is clear when we compare Fig. 6(c) with Fig. 6(d). When the zero-inflation mixing coefficient *ω*_*j*_ = 0, our transformation is

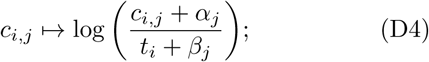

when *ω* ≠ 0, it is

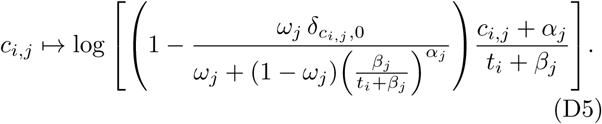

We provide two choices regarding how to compute the relative size factor *t*_*i*_ for the *i*-th cell: the first definition is

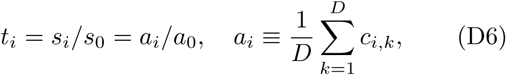

where the ratio *s*_*i*_/*s*_0_ is the same as that used in the log normalization method (D1), *a*_*i*_ is the arithmetic mean of the *D* surface protein counts, and *a*_0_ = *s*_0_/*D* is some choice of typical value of *a*_*i*_; the other definition is

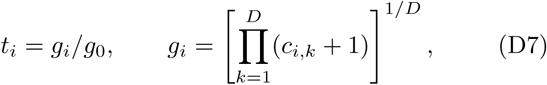

where *g*_*i*_ is the geometric mean as in the two versions of CLR transformation (D2) and (D3), and *g*_0_ is some choice of typical value of *g*_*i*_. Although (D6) capturing the differences in total count might seem more intuitive, we recommend (D7), as the geometric mean is more robust against outliers than the arithmetic mean. Transformation results for the two choices are shown in Fig. 7(d) and (e), respectively, where it is apparent that the second convention *t*_*i*_ = *g*_*i*_/*g*_0_ better separates the CD8+ T cells from spiked-in mouse cells along the CD3+ direction. This phenomenon may be attributed to the fact that CD8+ T cells have high CD8 counts and, thus, inflated size factors under the former definition, thereby suppressing the transformed CD3 values. Consistent with this observation, performing statistical tests on the CD3 level shows that the CD8+ T cells are correctly classified as being CD3+, when (D7), but not when (D6), is used to calculate the size factors.

In each of the transformation methods described above, the argument of the logarithm can be considered as a normalized version of the raw count *c*_*i,j*_. In our transformation, the argument is E[λ], and the normalization adds a data-driven pseudocount *α*_*j*_ to the raw count and *β*_*j*_ to the relative size factor *t*_*i*_ ~ 1, which corrects the sum *c*_*i,j*_ + *α*_*j*_ for different sequencing depths. Similarly, the term containing *ω*_*j*_ corrects for the case of an inflated zero count. Compared with the log normalization (D1) and the CLR transformation (D2,D3), our approach utilizes the model parameters inferred from the data, rather than adding an arbitrary pseudocount of 1. It is also specific to a particular surface protein and has the ability to address a potential dropout effect in the measurement.

In our implementation, we have chosen *g*_0_ to be the geometric mean of all *g*_*i*’_s. However, a different choice of *g*_0_ would merely translate the distribution of the transformed data by a constant. That is, under rescaling *t*_*i*_ → *bt*_*i*_ by a fixed constant *b*, the mixture probabilities of the NB and ZINB models remain invariant under the compensating redefinitions λ_*i,j*_ → λ_*i,j*_/*b*, *α*_*j*_ → *α*_*j*_, *β*_*j*_ → *bβ*_*j*_, (*c*_*i,j*_ +*α*_*j*_)/(*t*_*i*_ +*β*_*j*_) → (1/*b*)(*c*_*i,j*_ +*α*_*j*_)/(*t*_*i*_ +*β*_*j*_), and *ω*_*j*_ → *ω*_*j*_; hence, the only effect of choosing a different *g*_0_ is to rescale E[λ_*i,j*_] by a multiplicative constant for all *i*, resulting in translating log E[λ_*i,j*_] by a global constant for all cells.

